# Bacteriophage anti-defense genes that neutralize TIR and STING immune responses

**DOI:** 10.1101/2022.06.09.495361

**Authors:** Peiyin Ho, Yibu Chen, Subarna Biswas, Ethan Canfield, Douglas E. Feldman

## Abstract

Programmed cell suicide of infected bacteria, known as abortive infection (Abi), serves as a central immune defense strategy to prevent the spread of bacteriophage viruses and other invasive genetic elements across a population. Many Abi systems utilize bespoke cyclic nucleotide immune messengers generated upon infection to rapidly mobilize cognate death effectors. Here, we identify a large family of bacteriophage nucleotidyltransferases (NTases) that synthesize competitor cyclic dinucleotide (CDN) ligands, inhibiting NAD-depleting TIR effectors activated by a linked STING CDN sensor domain. Virus NTase genes are positioned within genomic regions containing other anti-defense genes, and through a functional screen, we uncover candidate anti-TIR defense (Atd) genes that confer protection against TIR-STING cytotoxicity. We show that a virus MazG-like nucleotide pyrophosphatase identified in the screen, Atd1, depletes the starvation alarmone (p)ppGpp, revealing a role for the alarmone-activated host toxin MazF as a key executioner of TIR-directed abortive infection. Phage NTases and counter-defenses like Atd1 preserve host viability to ensure virus propagation, and may be exploited as tools to modulate TIR and STING immune responses.

## Introduction

Bacteria are under the constant threat of attack from viruses (phages) and other mobile genetic elements (MGE), and have evolved a multitude of defenses, ranging from restriction enzymes to CRISPR/Cas adaptive immunity, to mitigate this threat. Programmed bacterial cell suicide, known as abortive infection (Abi), represents a primary strategy employed by infected cells to block the colony-wide propagation of viruses and achieve population-level immunity, at the expense of individual survival (Chopin et al., 2005; Lopatina et al., 2020).

Early studies on operon-linked toxin-antitoxin (TA) systems, as well as more recent discoveries on retrons, have exposed the regulatory logic underlying Abi immune responses. In contrast to restriction enzymes and CRISPR/Cas, which are tonically active but narrowly targeted to specific sequences in foreign, but not host, nucleic acids (Haurwitz et al., 2010; Jinek et al., 2012; Pingoud et al., 2005), highly cytotoxic Abi systems must first be activated by sensing intrusion of the MGE into the cell. TA systems are armed through a combination of a labile antitoxin and the convergence of multiple stressors, such as phage infection, that downregulate transcription of the TA operon, resulting in rapid loss of the antitoxin and activation of a lethal toxin (Harms et al., 2018; Jurėnas et al., 2022). Retrons are likewise held in an inactive conformation until their toxicity is unleashed through engagement with specific phage proteins (Millman et al., 2020).

Nucleotide-based immune alarm signaling has emerged as a central mechanism across distinct Abi systems. Type III CRISPR-Cas, Pycsar, and cyclic dinucleotide-based antiphage signaling systems (CBASS) are triggered through activation of constituent nucleotidyltransferases (NTases) and nucleotide cyclases upon phage infection (Kazlauskiene et al., 2017; Tal et al., 2021; Whiteley et al., 2019). Cyclic nucleotide messengers generated by these enzymes mobilize cognate death effectors, including nucleases, proteases, pore-forming toxins, as well as sirtuin and Toll-interleukin-1 receptor (TIR) domain NAD-cleaving enzymes (Kazlauskiene et al., 2017; Lowey et al., 2020; Ofir et al., 2021), through direct binding to an effector-linked cyclic nucleotide sensor, such as the widespread STING (stimulator of interferon genes) domain. Many viruses, in turn, deploy specialized nucleases that rapidly cleave and inactivate cyclic nucleotides, extinguishing host immune responses (Athukoralage et al., 2020; Hobbs et al., 2022). Other virus-encoded proteins act as molecular ‘sponges’ that sequester nucleotide immune messengers (Leavitt et al., 2022).

Currently, our understanding of how viruses evade the multitude of known Abi systems is limited, and little is known about whether viruses employ counter-defense strategies beyond signal degradation or interception. This stands in contrast to the large repertoire of experimentally characterized anti-CRISPRs, many of which inactivate CRISPR/Cas subtypes with exquisite specificity (Bondy-Denomy et al., 2015; Marino et al., 2018). Here, we uncover a family of virus-encoded NTases that generate CDN competitor ligands that impair activation of TIR death effectors containing a linked STING CDN sensor domain (TIR-STING). We show that virus genes encoding NTases are positioned in close proximity to known anti-defense genes, and through a plasmid-based functional screen, we identify novel virus anti-TIR defense (Atd) candidate genes. Among these, we uncover a phage MazG-like nucleotide pyrophosphatase, Atd1, that depletes the starvation alarmones (p)ppGpp, revealing how TIR-mediated abortive infection is executed, in part, through alarmone-activated host toxins.

## Results

Bioinformatic analysis of bacteriophage genomes across the *Siphoviridae, Myoviridae* and *Podoviridae* virus families revealed a large group of genes encoding predicted NTases that are closely related to the minimal NTases (MNT) of the polβ NTase superfamily (4), with a more distant relationship to kanamycin NTases (KNT) (Figure 1A). While phage NTases contain an intact catalytic site GS-x-AY[GAN]T-x4-SDxD that is similar to that found in MNT, they lack a near-invariant tyrosine (Y) residue that is positioned immediately C-terminal to the GS motif in MNT (Figure 1B, site 1). Furthermore, phage NTases, but not MNT, contain a conserved NP-x-h2[DE] sequence positioned approximately 60 residues C-terminal to the SDxD motif (Fig, 1B, site 2). This sequence divergence supports the proposal that phage NTases encompass a distinct clade within the NTase superfamily.

**Figure 1.**
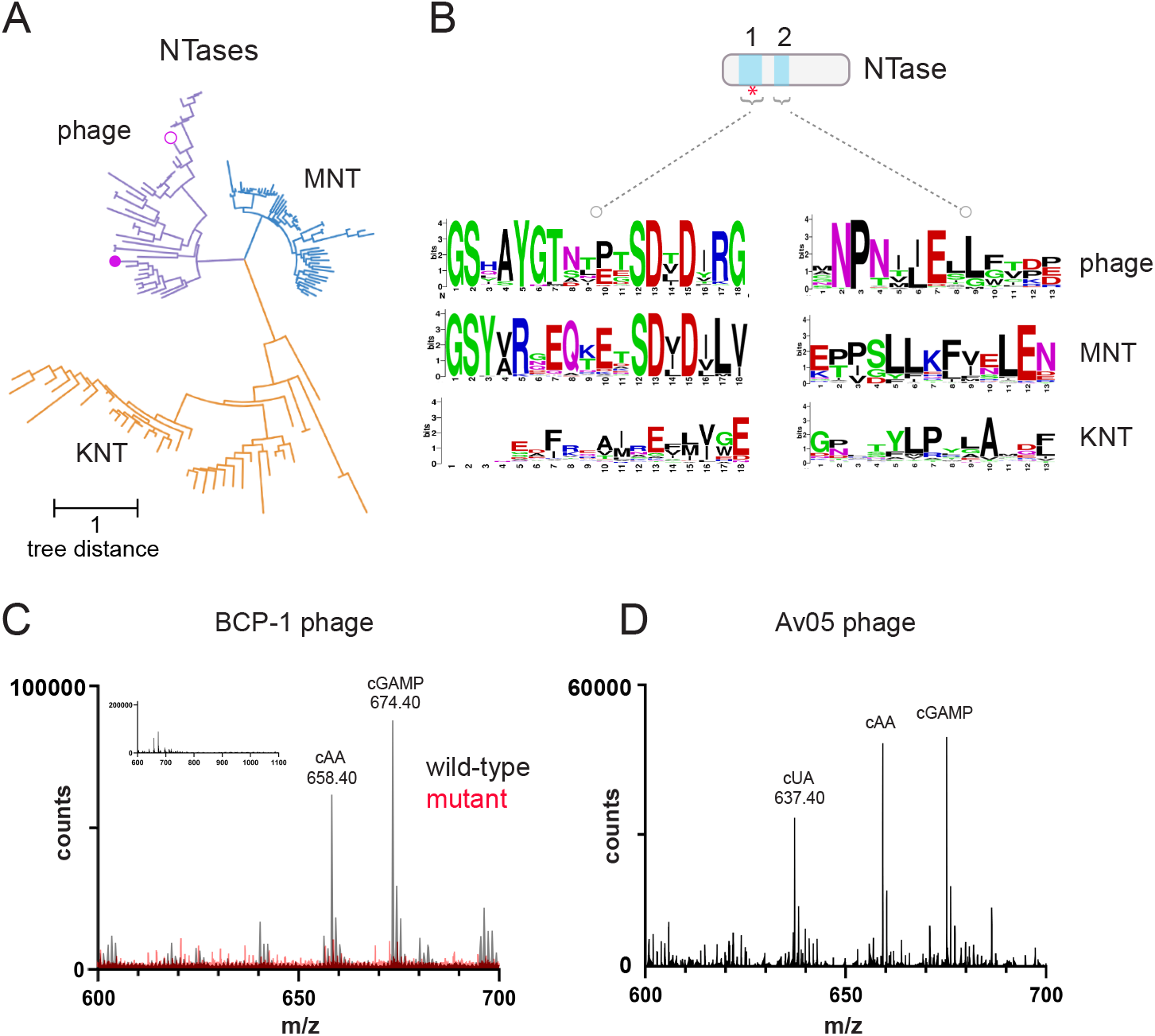
Phage nucleotidytransferases synthesize CDN products. (A) Dendrogram of phage NTases, MNT, and KNT clades within the DNA polymerase-β superfamily, built using alignment of protein sequences. Open and filled purple circles respectively indicate Av05 and BCP1 phage NTases analyzed in this study. (B) Sequence LOGO diagram showing divergence of NTase sequences across two sites. Red asterisk indicates major catalytic site. (C) Overlaid MALDI-MS chromatograms showing products generated by wild-type and mutant BCP1 phage NTase. Major products with masses corresponding to 3’3’cGAMP and cAA are shown. Inset shows chromatogram for wild-type BCP1 NTase across a wider m/z range. Results are representative of n=3 experiments. (D) MALDI-MS chromatogram showing reaction products generated by Av05 phage NTase. Results are representative of n=2 experiments. See also Figure S1.

To examine possible enzymatic functions of phage NTases, we incubated purified, recombinant NTases from the *Bacillus* phage BCP1 and Enterobacteriophage Av05 in the presence of rNTPs. Wild-type BCP1 NTase, but not a catalytic site mutant (SDWD→AAWA), generated 3’3’cGAMP and cyclic di-adenylate (cAA) as major products, as determined by MALDI-MS profiling (Figures 1C, S1A and S1B). Av05 NTase likewise generated 3’3cGAMP and cAA, as well as a product consistent with the mass of cyclic uridine-adenylate (cUA) (Figures 1D and S1C). These findings expand the range of NTase families that are capable of synthesizing CDN products.

While 3’3’cGAMP acts as an immune alarm signal that mobilizes host CBASS death effectors (Cohen et al., 2019; Lowey et al., 2020; Whiteley et al., 2019), the widespread distribution of NTases across virus families suggests that phage CDN synthesis may confer a fitness advantage. In line with this proposal, 3’3’cGAMP binds, with varying affinities, to STING CDN sensor domains contained within TIR-STING effectors from different species, and can act as a competitive inhibitor of the activating ligand cyclic di-guanylate (cGG) (Morehouse et al., 2020). We therefore sought to explore the effect of phage NTases on TIR-STING, using cleavage of the fluorogenic substrate ε-NAD as a measure of enzymatic activity. Co-incubation of *S. falciparum* (*Sf*) TIR-STING with cGG and increasing amounts of BCP1 and Av05 NTases resulted in a concentration-dependent inhibition of the initial ε-NAD cleavage rate, while a BCP1 catalytic mutant had no effect on TIR-STING NADase activity (Figure 2A). When introduced into *E. coli*, BCP1 and Av05 phage NTases protected hosts against the toxic effects of *Sf*TIR-STING, as determined through colony forming unit (CFU) assays, while the BCP1 catalytic mutant failed to do so (Figure 2B). In contrast, BCP1 NTase was largely ineffective in inhibiting the enzymatic activity of *Capnocytophage granulosa* (*Cg*) TIR-STING, which exhibits a 100-fold lower affinity for 3’3’cGAMP relative to that of *Sf*STIR-STING (Morehouse et al., 2020) (Figure S2A), and did not protect hosts against *Cg*TIR-STING (Figure S2B). Collectively, these results suggest that phage NTases generate competitor CDN ligands such as 3’3’cGAMP as a strategy to inhibit TIR-STING immune effectors.

**Figure 2.**
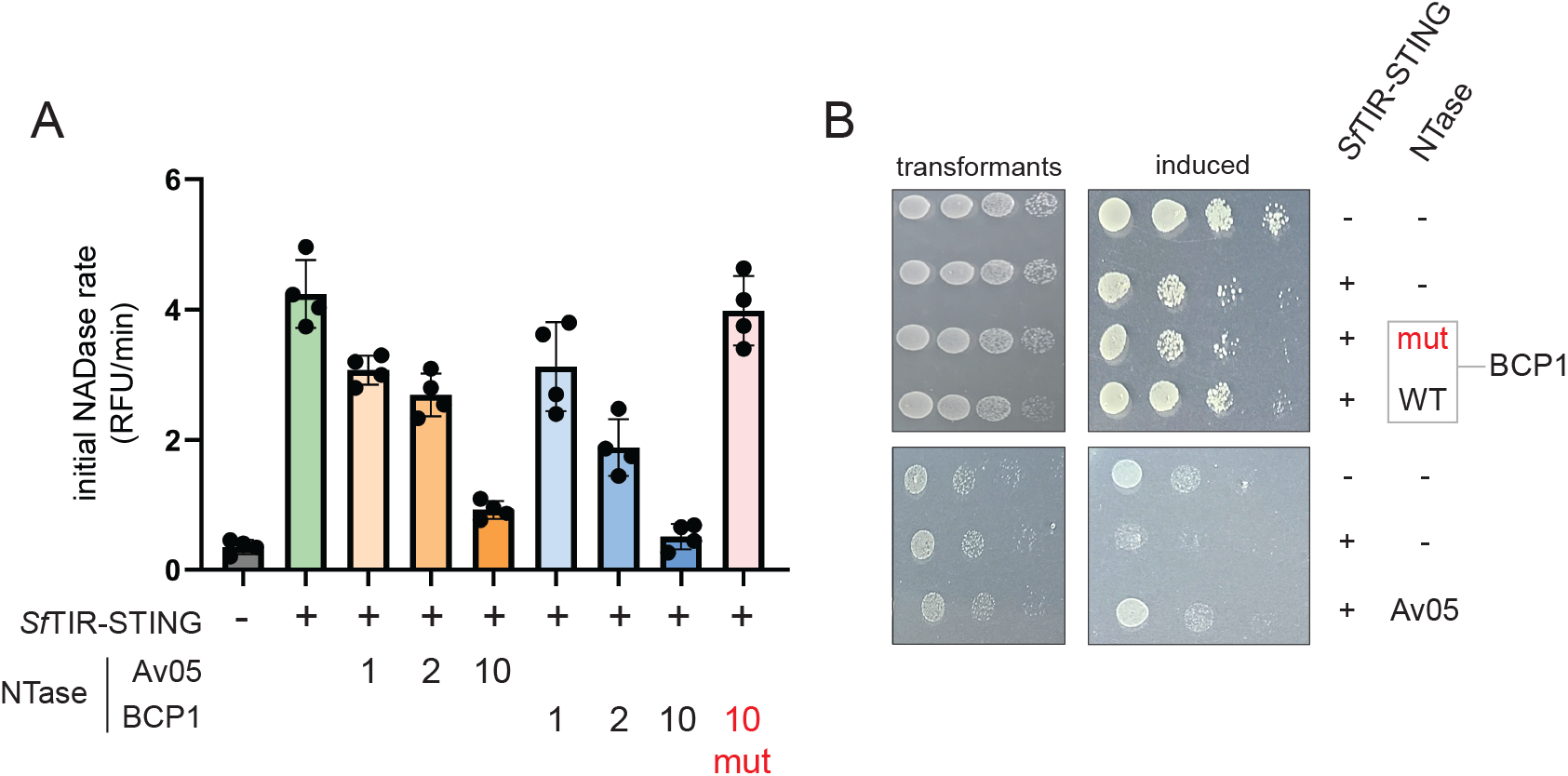
Phage nucleotidyltransferases impair the activity of a TIR-STING death effector. (A) NAD+ hydrolysis assay. *S. falciparum (Sf*) TIR-STING NAD+ cleavage activity was measured using the fluorescent substrate ε-NAD, in the absence or presence of increasing amounts of wild-type or catalytically inactive forms of BCP1 and Av05 phage NTases across a 10-fold range in concentration. Graphs show mean values with s.d. from n=4 experiments. (B) CFU assay showing relative viability of *E. coli* transformed with the indicated combinations of SfTIR-STING and the wild-type or catalytic mutant forms of BCP1 or Av05 NTase, or vector control. Results are representative of n=3 experiments. See also Figure S2.

Phage anti-restriction and anti-CRISPR genes are often positioned together within larger anti-defense gene clusters (Marino et al., 2018; Pinilla-Redondo et al., 2020). We therefore performed a bioinformatics analysis of genes flanking phage NTases. Starting from a set of 240 phage NTases (Table S1), we identified 1958 flanking open reading frames (ORFs), encompassing 332 clusters of closely related sequences. Among these, we found DNA adenine and cytosine methyltransferases, which block host restriction endonucleases, and NAD+ biosynthetic enzymes that may mitigate the metabolite-depleting effects of TIR NADases (Figure 3A). This suggested that previously unrecognized anti-defense genes may be positioned in close proximity to genes encoding NTases.

**Figure 3.**
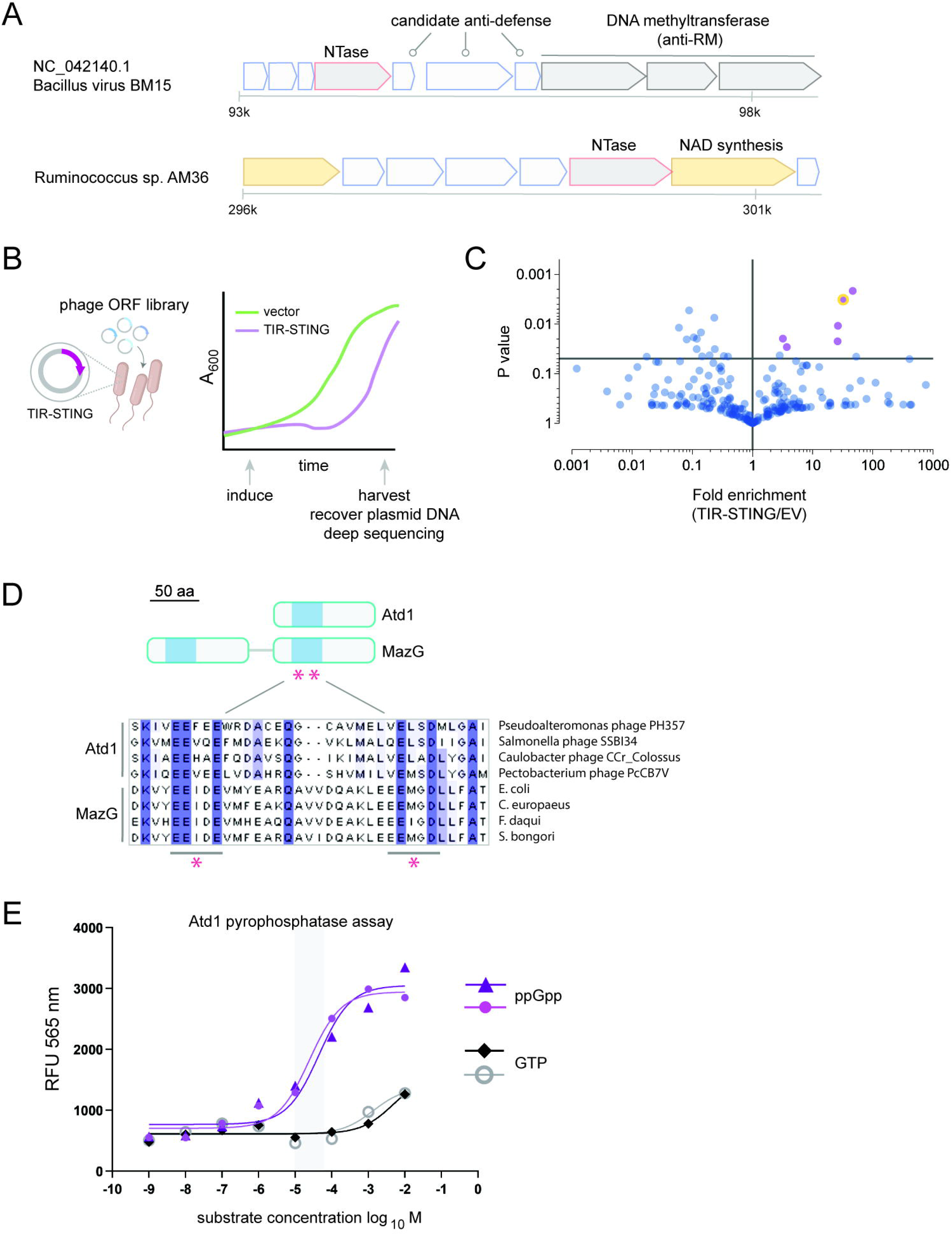
Functional screening identifies Atd1, a phage MazG-like nucleotide pyrophosphatase that confers protection against TIR-STING toxicity. (A) Examples of phage genomic loci containing NTases in close proximity to anti-restriction DNA methyltransferases and NAD biosynthetic enzymes. (B) Schematic of plasmid-based functional screen. A plasmid library of phage genes selected on the basis of genomic proximity to NTase-encoding genes was transformed into *E. coli* hosts together with empty vector (EV) or inducible TIR-STING plasmid. Culture samples were withdrawn immediately prior to induction or following overnight growth. The relative representation of genes within the phage library for each strain and at each time point was then determined by deep sequencing. (C) Volcano plot showing mean enrichment of phage genes in host cells co-expressing TIR-STING, normalized to EV controls. Phage genes significantly enriched (*P* < 0.05) in hosts co-expressing TIR-STING are displayed in purple. Atd1 is indicated with an orange outline. (D) Top, schematic representation of Atd1 and MazG proteins. The twin catalytic motifs are denoted by red asterisks. Bottom, multiple sequence alignment of phage and bacteria MazG family members. Conserved catalytic residues are highlighted in blue. (E) Pyrophosphatase assay measuring activity of Atd1 on ppGpp and GTP substrates across a range of concentrations. The shaded area shows the 95% confidence interval of the Km of Atd1 for ppGpp. Results from two independent replicate experiments are plotted for each substrate. See also Figure S3.

To identify additional candidate anti-defense genes, we performed a plasmid-based functional screen for phage ORFs that are selectively enriched when co-expressed with a TIR-STING death effector, but not in the presence of an empty vector control. We first co-transformed *E. coli* with an inducible phage ORF library encompassing 257 NTase-adjacent ORFs, representative of 176 sequence clusters, together with a second inducible plasmid encoding TIR-STING or an empty vector control. Each transformant strain was then inoculated into liquid culture, and subsequently shifted to growth under inducing conditions before cell recovery (Figure 3B). We PCR amplified the phage insert region of the input plasmid library and the output plasmid library following overnight growth under inducing conditions, and used next-generation sequencing to compare the representation of individual inserts in the input and output plasmid libraries.

Following this approach, we identified six phage genes that are significantly enriched in bacterial hosts co-expressing TIR-STING, but not an empty control plasmid, suggesting that these genes confer protection against TIR-STING toxicity and thus provide a survival advantage (Figure 3C). While most ORFs showed no detectable sequence similarity to any previously characterized protein, a gene from *Pectobacterium* phage PcCB7V encoding the catalytic domain of a MazG-like nucleotide pyrophosphatase was significantly enriched in the screen (Figures 3C and 3D) (Lee et al., 2008). In *E. coli, mazG* is positioned downstream of *relA* within the *mazEF* operon, a stress-induced TA suicide module. MazF is a stable endoribonuclease toxin that is inhibited through a direct interaction with its cognate antitoxin MazE, a labile protein (Gross et al., 2006). Under conditions of nutrient deprivation, RelA senses deacylated tRNA in the ribosomal A-site of starved ribosomes, and synthesizes the alarmone nucleotides guanosine pentaphosphate (pppGpp) and tetraphosphate (ppGpp)—collectively referred to as (p)ppGpp—that globally downregulate transcription as part of the stringent response, resulting in rapid loss of the MazE antitoxin and activation of the promiscuous MazF endoribonuclease toxin (Culviner and Laub, 2018; Engelberg-Kulka et al., 2005).

The identification of a phage MazG-like protein, which we have provisionally designated Atd1 (*a*nti-*T*IR *d*efense 1), as a hit in the screen suggested that enzymatic depletion of (p)ppGpp may attenuate TIR-mediated cell suicide. Consistent with this possibility, purified wild-type Atd1 stimulated the dephosphorylation of ppGpp substrate with an apparent Km of ∼50 µM (Figures 3E and S3A), a value that is above intracellular concentrations of (p)ppGpp during logarithmic growth, but would allow for maximal enzyme activity following induction of the stringent response (Kriel et al., 2012; Varik et al., 2017). In contrast, no effect of Atd1 was seen on dephosphorylation of rNTPs or cGG under the same reaction conditions (Figures 3E and S3B).

We next utilized an RNA-based fluorescent (p)ppGpp sensor (Sun et al., 2021) to monitor the effects of TIR-STING expression on cellular alarmone pools. Live-cell imaging revealed significant fluorescence activation following induction of TIR-STING (Figure 4A), with a mean 2.7-fold increase in signal intensity relative to glucose-fed control cells, similar to the increase seen in cells exposed to a chemical inducer of starvation, α-methylglucose (α-MG) (Figures 4A and 4B). Conversely, phage MazG suppressed reporter fluorescence following 1 h exposure to α-MG (Figures 4C and 4D). Taken together, these observations indicate that TIR-STING triggers a rapid increase in cellular (p)ppGpp pools, and that this response can be suppressed by Atd1.

**Figure 4.**
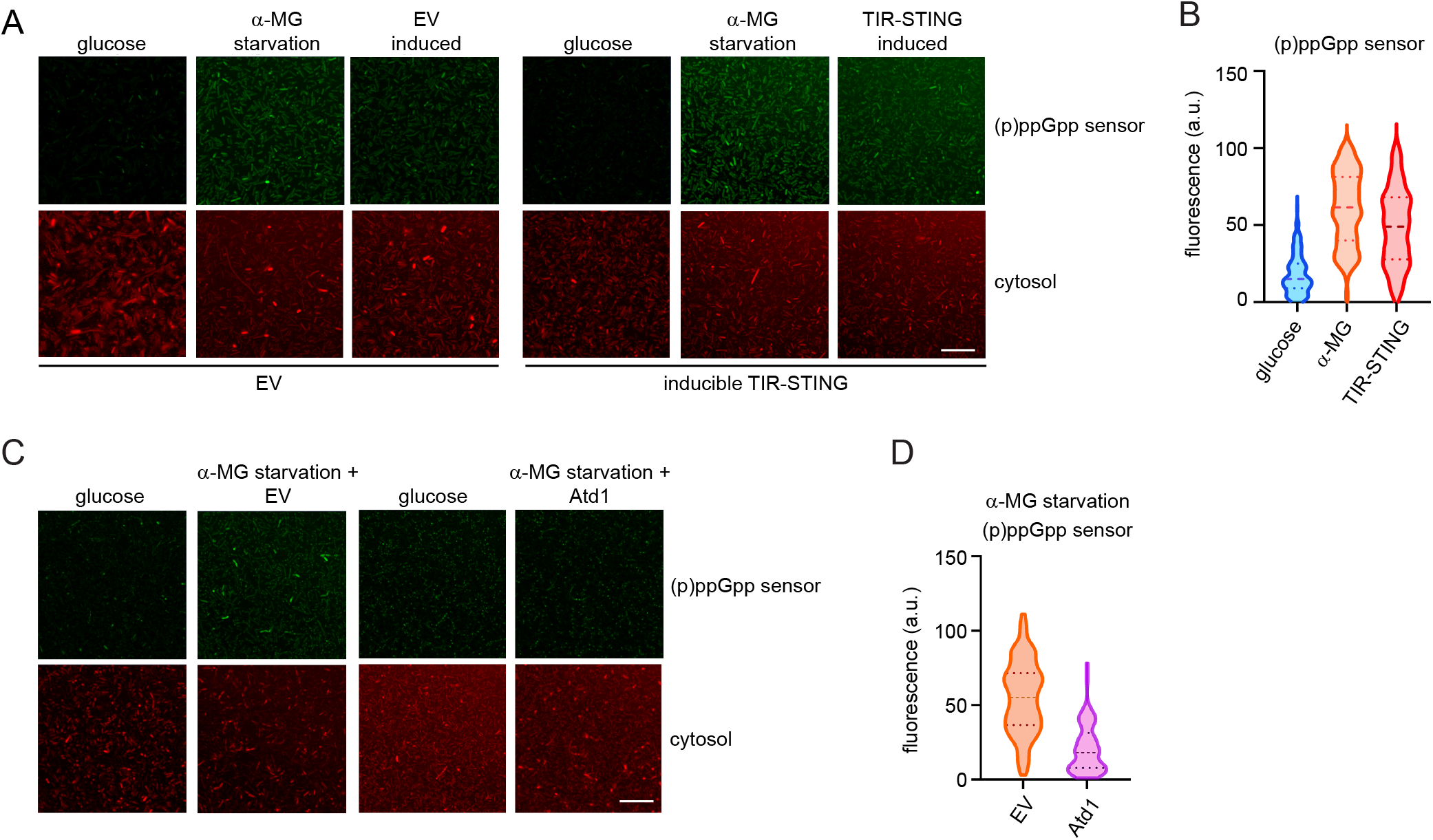
Atd1 depletes the starvation alarmone (p)ppGpp. (A) Visualization of cellular (p)ppGpp levels using RNA-based fluorescent sensor (green) and cytosolic RFP (red) in live bacteria grown in media containing glucose, or following treatment with the glycolysis inhibitor α-MG to induce starvation, or following induction of TIR-STING or control vector. Scale bar, 50 µm. Results are representative of n=3 experiments. (B) Violin plot of (p)ppGpp sensor signal intensities in live cells grown in glucose, treated with the chemical starvation agent α-MG, or following induction of TIR-STING. Dashed lines indicate quartiles. (C) Cellular (p)ppGpp sensor (green) and cytosolic RFP fluorescence (red) in cells grown in complete media containing glucose, or following starvation induced by α-MG in the absence or presence of Atd1. Scale bar, 50 µm. Results are representative of n=3 experiments. (D) Violin plot of (p)ppGpp sensor signal intensities in live cells following treatment with α-MG, in the presence of Atd1 or empty control vector. Dashed lines indicate quartiles.

To assess the contribution of (p)ppGpp and the stringent response to TIR-STING toxicity, we performed CFU assays to measure and compare the viability of *mazG, relA* and *mazEF* mutants expressing TIR-STING, relative to wild-type parental control strains. Loss of *mazG*, which confers hypersensitivity to amino acid starvation (Gross et al., 2006), led to a 4-fold further decrease in viability upon TIR-STING expression, relative to a parental strain (Figures 5A and 5B). Conversely, mutational inactivation of *relA* or *mazEF* protected cell viability from TIR-STING toxicity (Figures 5A and 5B). Bacteriophage lambda RexB, which inhibits the proteolytic degradation of MazE to maintain the MazF toxin in an inactive conformation (Engelberg-Kulka et al., 1998), also increased the viability of cells co-expressing TIR-STING, as did Atd1, validating the results from the primary screen (Figure 5A and 5B).

**Figure 5.**
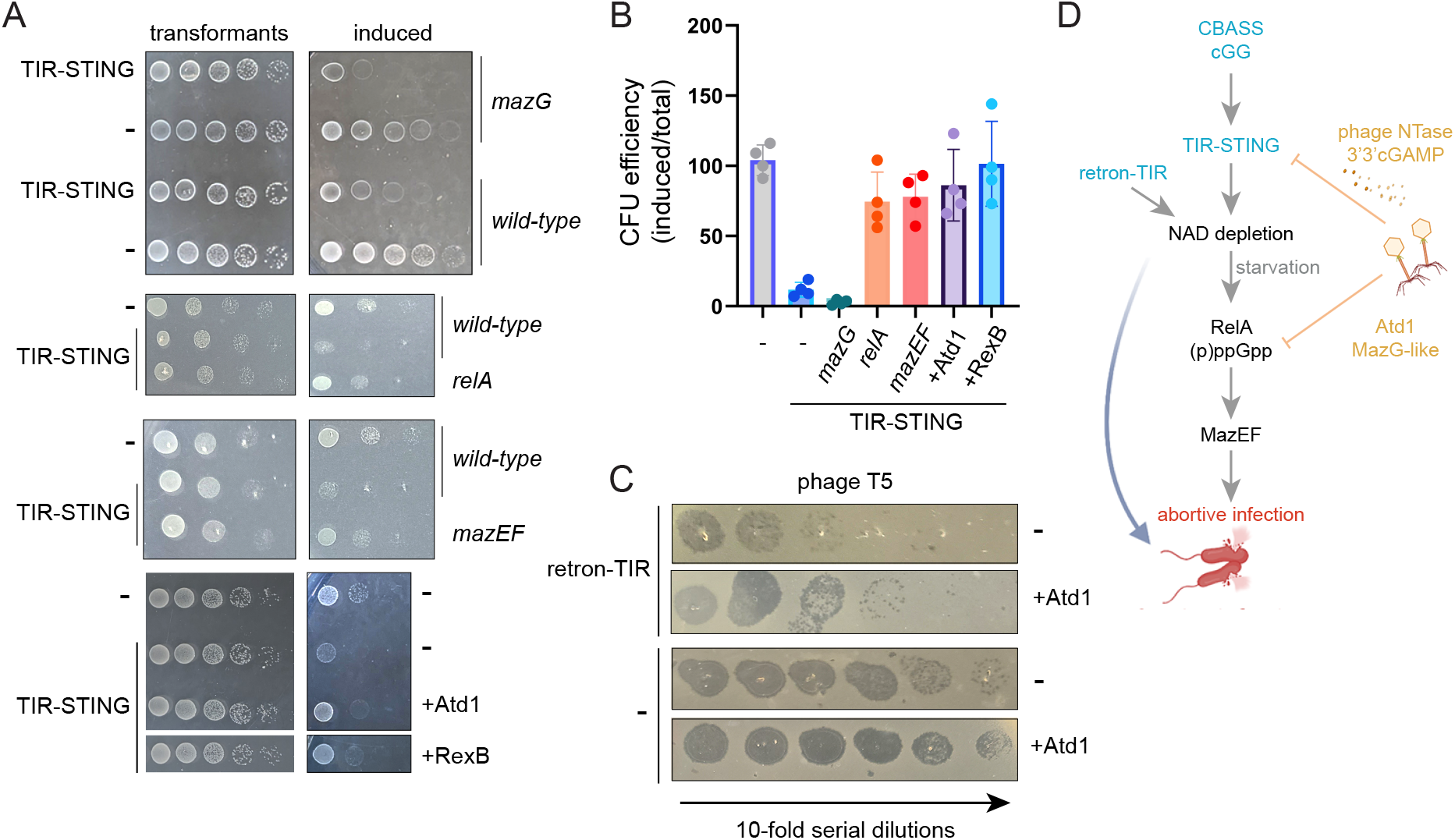
Role of RelA and the MazF toxin in promoting TIR-dependent abortive infection. (A) CFU assays showing effects of genetic inactivation of *mazG, relA* or *mazEF*, or expression of RexB or Atd1, on TIR-STING cytotoxicity. Results are representative of n=4 replicate experiments. (B) Quantification of CFU assay results. Bar graphs show mean values, error bars show s.d. from n=4 experiments. (C) Phage plaque formation assay. Bacterial lawns grown from *E. coli* expressing a retron-TIR defense system or carrying an empty vector control, each in the absence or presence of Atd1, were infected with droplets containing 10-fold serial dilutions of T5 phage. Results are representative of n=3 replicate experiments. (D) Model depicting suppression of host immunity by phage NTases and Atd1, which intercede at distinct points in abortive infection responses orchestrated by immune effectors containing TIR and STING domains.

Notably, Atd1 also blunted the toxic effects of a distinct TIR domain death effector, retron-TIR (Gao et al., 2020), rescuing by 10 fold plaque formation by the T5 bacteriophage, relative to a control strain that did not express Atd1 (Figure 5C). These observations together indicate that Atd1 can confer protection against distinct defense systems that utilize TIR domain-containing effectors (Figure 5D).

## Discussion

Abortive infection is an immune strategy predicated on host self-destruction and is designed to rapidly extinguish the spread of viruses and other invasive genetic elements across a bacterial population, at the expense of individual cell survival (Lopatina et al., 2020). Here, we uncover virus anti-defense genes that counteract cytotoxic Abi effectors containing the widespread and highly conserved TIR and STING domains. These anti-Abi systems operate through distinct, nucleotide-centered mechanisms: the production of competitive inhibitor CDNs that can “jam” STING receptors, and the enzymatic depletion of (p)ppGpp alarmone molecules, impairing the host response to metabolite depletion by NAD-hydrolyzing TIR enzymes and the ensuing mobilization of cytotoxins.

We identify a large family of virus-encoded NTases that synthesize cAA and 3’3’cGAMP as major products, the same CDNs that act as immune messengers generated by bacterial CD-NTases encoded within host CBASS defense systems (Whiteley et al., 2019). Virus NTases are closely related to bacterial and archaeal MNT within the polβ-type NTase superfamily, while exhibiting a more distant sequence relationship to kanamycin NTases, which confer resistance to aminoglycoside antibiotics (Pedersen et al., 1995). These observations expand the scope of NTase families known to synthesize CDNs, and suggest that bacterial MNT enzymes may likewise generate CDN products. In contrast to CD-NTases and nucleotide cyclases contained, respectively, within CBASS and Pycsar systems (Cohen et al., 2019; Tal et al., 2021; Whiteley et al., 2019), phage NTase are not encoded within operons containing known cyclic nucleotide sensor or effector domains. Rather, by generating 3’3’cGAMP, virus NTases may competitively inhibit host STING receptors, many of which can bind to distinct CDNs but are selectively activated by cGG (Morehouse et al., 2020). It will be important to determine whether this signal jamming strategy extends beyond TIR-STING to other classes of nucleotide-sensing effectors, such as those containing the widespread SAVED, patatin and SLOG domains (Duncan-Lowey et al., 2021; Lowey et al., 2020; Ofir et al., 2021). Our results additionally reveal the general design for a second virus anti-suicide system that exploits a previously unrecognized dependency of TIR NADase-driven cell death on production of the stringent response alarmones (p)ppGpp. We show that a virus MazG-like protein, Atd1, depletes cellular (p)ppGpp pools, blocking the mobilization of host TA suicide modules, including MazEF. Thus, while depletion of the essential metabolites NAD+ and NADP is thought to induce metabolic arrest (Morehouse et al., 2020), the combinatorial engagement of secondary death pathways may be required for maximal toxicity of TIR effectors. The transmission of cytotoxic signals downstream of TIR may occur not only through the activation of host metabolic stress pathways, as shown here, but additionally through the generation of the NAD cleavage product cyclic ADP ribose (cADPR), which itself can act as a messenger that mobilizes downstream immune effectors (Essuman et al., 2017; Essuman et al., 2018; Ofir et al., 2021).

The identification of anti-Abi gene neighborhoods within phage genomes opens avenues for the systematic screening and identification of new classes of immune modulators. The diversity of bacterial Abi defenses, and the conservation and modularity of key effector domains such as TIR and STING (Gao et al., 2020; Ofir et al., 2021), highlights the potential for anti-Abi systems in the engineered control of immune signaling and cell survival.

## Methods

### Evolutionary and LOGO sequence analysis of NTase enzymes

Bacteriophage genes within the *Siphoviridae, Myoviridae* and *Podoviridae* families annotated as nucleotidyltransferases and containing intact catalytic motifs were used as query sequences for iterative BLASTp searches. Evolutionary analyses were conducted with MEGA software (Tamura et al., 2021) using the Maximum Likelihood method and JTT matrix-based model (Jones et al., 1992). The tree with the highest log likelihood is shown. Initial trees for the heuristic search were obtained automatically by applying Neighbor-Join and BioNJ algorithms to a matrix of pairwise distances estimated using the JTT model, and then selecting the topology with superior log likelihood value. Trees were visualized using iTOL v6 (Letunic and Bork, 2021). The tree is drawn to scale, with branch lengths measured in the number of substitutions per site. LOGO sequence analysis of NTase catalytic domains was performed using WebLogo (Crooks et al., 2004).

### Recombinant protein expression and purification

Synthetic gene fragments (gBlocks, IDT) encoding BCP1 and Av05 phage NTases, *Sf* and *Cg* TIR-STING, and Atd1 and containing an N-terminal TEV cleavage site were cloned directly via Gibson assembly (NEBuilder HiFi, New England Biolabs) into BamHI/NotI-digested pGEX4-1 (GE Healthcare), or tagged with N-terminal TwinStrep (TS) or streptavidin binding peptide (SBP) tags by PCR and introduced into NdeI/XhoI-digested pET30a vector (MilliporeSigma). Mutagenesis was performed using the Q5 Site-Directed Mutagenesis kit (New England Biolab). All constructs were confirmed by Sanger sequencing on both strands. Plasmids were transformed into BL21 (DE3) Rosetta 2 competent cells (MilliporeSigma). Bacterial cultures were grown to an A_600_ of 0.4-0.5, then induced with 0.1 mM IPTG for 16 hours at 30°C. Bacterial pellets were resuspended in ice-cold PBS supplemented with complete protease inhibitor cocktail, 5 mM sodium fluoride, 1 mM sodium orthovanadate, lysozyme (0.2 mg/ml) and DNase I (10 μg/ml), then sonicated in 5-6 bursts of 15 s with 30 s cooling in between bursts. Soluble proteins (5 ml) were incubated with 250 μl magnetic glutathione resin (ThermoFisher) or streptactin resin (IBA Life Sciences) for 2 h at 4°C with gentle rotation. Resin was washed three times in lysis buffer and eluted twice for 30 min in elution buffer (50 mM Tris-HCl pH 8.0, 200 mM NaCl, 5% glycerol, 10 mM desthiobiotin or glutathione), then pooled and dialyzed overnight at 4°C into TBS containing 5% glycerol. Samples were aliquoted and stored at -80°C.

### Nucleotide synthesis assays

In vitro biochemical reactions were performed with 1 uM purified enzyme in NTase reaction buffer (50 mM Tris pH 8.0, 50 mM KCl, 5 mM Mg(OAc)_2_, 1 mM DTT, 5% glycerol, 125 μM rNTPs) in a final volume of 20 µl for 3 h at 37°C. Reactions were deproteinized for MALDI-MS analysis with 80 µl acetonitrile and stored at -20°C prior to analysis.

### MALDI-MS analysis

Samples were desalted and concentrated using ZipTip C18 micropipette tips (MilliporeSigma), then spotted on a 96-spot steel plate target and covered with alpha-cyano-4-hydroxy-cinnamic acid (HCCA) matrix solution. Mass spectrometry analysis was performed with a Microflex MALDI-TOF mass spectrometer (Bruker Daltonics) and spectra were recorded for positive ions. Polytools (2.0) software was used to analyze mass spectra.

### TIR NADase assay

In vitro TIR NADase assays were performed essentially as described. Briefly, purified TIR-STING (0.5 µM) was incubated in TIR reaction buffer (20 mM HEPES-KOH pH 7.5, 100 mM KCl) containing 0.25 mM ε-NAD substrate and supplemented with 125 μM rNTPs, in the absence or presence of increasing amounts of purified Av05 NTase or the wild-type or mutant forms of BCP1 NTase 0.125, 0.25,1.25 µM). Reactions were performed in 96-well plates and initiated through the addition of cGG (40 nM), and read continuously using Synergy H1 Hybrid Multi-Mode Reader (BioTek) in fluorescence mode at 410 nm with excitation at 300 nm. Initial reaction rates are calculated using values obtained immediately prior to cGG addition, and at the 10 min time point.

### Bacterial CFU assays

*relA* and *mazG* mutant and wild-type parental *E. coli* strains were obtained from the Keio Strain Collection (Horizon Discovery). The *mazEF* mutant and parental strains were a generous gift from Dr. Hanna Engelberg-Kulka. Cells from overnight cultures were diluted 1:20 in 5 ml of M9 medium containing 0.25% casamino acids (CAA) and 0.5% glucose. Cultures were grown with shaking at 200 rpm at 37°C to early log phase (OD_600_ 0.2-0.3). Cells were then washed in M9 minimal medium, serially diluted, and plated on M9 agar plates containing 0.25% CAA and 0.25% glucose; or 1% glycerol, 0.7% arabinose, as well as 0.5 mM IPTG for pET vector induction. Plates were incubated at 37°C overnight, and cell survival was evaluated by comparing the colony-forming ability of cells under inducing and nutrient limiting conditions, normalized to the total number of viable cells grown on plates containing glucose.

### Computational analysis of phage NTase-adjacent ORFs

Phage NTases were compiled at NCBI Protein and aligned using Jalview (Waterhouse et al., 2009) to confirm the presence of intact catalytic motifs (GS and DXD). Protein-coding ORFs positioned within 4 genes of phage NTases were identified using webFlaGs (Saha et al., 2021) and their corresponding protein sequences downloaded using NCBI Batch Entrez. To reduce redundancy, highly similar sequences (at least 98% sequence identity and coverage) were discarded using the linclust option in MMseqs2 (Hauser et al., 2016; Mirdita et al., 2019) with parameters --min-seq-id 0.98 -c 0.98, and the resulting sequences clustered based on homology using the cascaded clustering option in MMSeqs2 to yield a list of 1603 ORFs distributed across 332 clusters. Uncharacterized ORFs less than 200 amino acids in length (257 ORFs representative of 176 clusters) were selected for functional screening studies.

### Phage ORF library screening

Double-stranded gene fragments and pooled, single-stranded DNA oligonucleotides encoding phage ORFs with 20-25 nt flanking regions of homology to the pET-SUMO2 vector flanking the NdeI and NotI sites were purchased from IDT (eBlocks and oPools). ssDNA oligos were converted to dsDNA through isotheral extension of a primer complementary to the 3’ flanking region (sparQ HiFi PCR Master Mix, Quantabio), and products purified with a DNA spin column. Pooled ORFs were inserted into pET-SUMO2 by DNA assembly (NEBuilder HiFi) and transformed into *E. coli* (NEB DH5α). Colonies were scraped from plates, combined and inoculated into 10 ml liquid cultures. Plasmid DNA encoding the pooled ORF library was then prepared and used to transform BL21 (DE3) Rosetta 2 competent cells (MilliporeSigma) together with pET30a encoding Flag-TIR-STING or empty vector control. 100 µl of overnight cultures were diluted 1:20 and grown in M9 media supplemented with 0.25% CAA and 1% glycerol, and induced at A_600_=0.3-0.4 through the addition of 0.5mM IPTG. Samples were taken immediately prior to induction and at 18 h post-induction. Plasmid DNA was recovered by miniprep, and library inserts amplified by PCR using primers flanking the ORF insertion site in pET-SUMO2. Bar-coded libraries for Illumina sequencing were generated using the sparQ DNA Library Kit (Quantabio) and 50 bp reads generated using a HiSeq3000 sequencing system.

### Pyrophosphatase assay

Phosphatase reactions were performed in 96-well plates in pyrophosphatase reaction buffer (50 mM Tris-HCl (pH 7.5), 250 mM KCl, 5 mM MgCl2, 1 mM TCEP) containing 1 µg of purified Atd1 protein and ppGpp, NTPs, or cGG for 1.5 h at 37°C. Reactions were stopped through addition of 10 µl of ice-cold 20 mM EDTA. Phosphate levels in each reaction were measured using the PiPer Pyrophosphate Assay Kit (ThermoFisher) following the manufacturer’s instructions. Absorbance values determined using a Synergy H1 Hybrid Plate Reader set for absorbance at 565 nm.

### Live cell imaging

*E. coli* BL21(DE3) cells carrying the S2 fluorescent sensor plasmid (Sun et al., 2021) were grown in M9 medium supplemented with 0.25% CAA and 0.5% glucose to an A_600_ of 0.2-0.4. Cells were briefly pelleted (8,000 x g, 45 s), then shifted to M9 medium containing 0.25% CAA, 0.5% glucose and 0.5mM IPTG, in the absence or presence of 2.5% α-MG to induce chemical starvation; or M9 containing 1% glycerol, 0.5mM IPTG and 0.75% arabinose to induce both nutrient deprivation and expression of TIR-STING from the pBAD vector. Cells were incubated with DFHBI-1T dye 1 h after shift to inducing media as described (Sun et al., 2021). Images were obtained using a Zeiss LSM 880 Airyscan confocal microscope and were quantified using ImageJ (1.53r) software.

### Phage plaque assay

Bacterial overnight cultures grown from a single colony of *E. coli* K-12 strain MG1655 that had been transformed with retron-TIR from *Shigella dysenteriae* NCTC2966 (Addgene #157883) or an empty vector control (pACYC184) were grown in LB media, and a 1:100 dilution grown in a shaker at 37ºC at 200 rpm until the A_600_ reached 0.4-0.5. Next, 3 ml pre-warmed top agar (M9 salts, 0.25% CAA, 1% glycerol, 0.75% agar) was mixed with 0.15 ml bacteria culture, and poured evenly onto agar plates containing 25 ug/ml chloramphenicol. T5 phage (ATCC 11303-B5) was propagated by the double agar overlay method, and plaques scraped into LB media. Phage samples were centrifuged to pellet cellular debris, and the supernatant was filtered through with a 0.22 µm sterile filter. Phage preparations were serially diluted in phage dilution buffer (20 mM Tris pH 8.0, 150 mM NaCl, 8 mM MgS0_4_) and 4 µl droplets of each dilution spotted onto the cooled top agar layer. Plaques were allowed to form for 16 h at 37°C.

## Acknowledgements

This project is supported in part by grant 5R21AA027535-02 from the NIH, IRG-16-181-57 from the American Cancer Society, and the USC Department of Pathology. We thank Dr. Alireza Abdolvahabi for assistance with MALDI-MS experiments and Dr. Aaron Whiteley for helpful discussions. We have no conflicts of interest to disclose.

## Author Contributions

P.H. performed biochemical assays and experiments with bacterial cells, and prepared libraries for next-generation sequencing. E.C. directed mass spectrometry analysis. Y.C. and S.B. performed bioinformatics analysis. P.H., and D.E.F. analyzed the data and wrote the manuscript.

