## Supplementary figures and images for "Bacteriophage anti-defense genes that neutralize TIR and STING immune responses"

### Figure S1

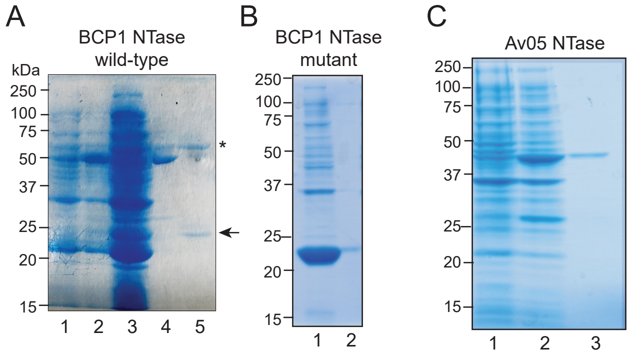

### Figure S2

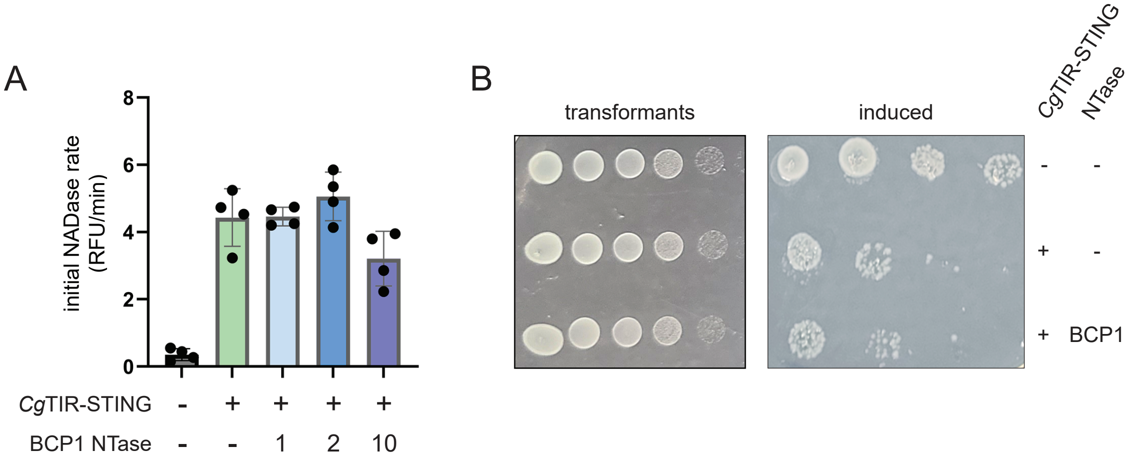

### Figure S2

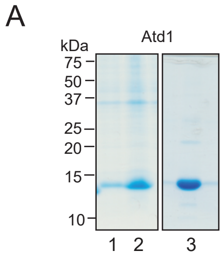

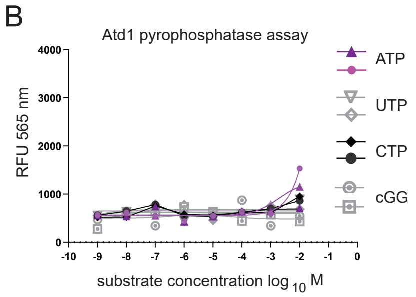
