## Supplementary material for "Bacteriophage anti-defense genes that neutralize TIR and STING immune responses": Supplemetary Figure Legends

**Figure S1. Purification of recombinant phage NTases, Related to Figure 1**

(A) Total protein was recovered from uninduced bacteria (lane 1) or postinduction cultures of bacteria expressing BCP1 NTase containing an N-terminal GST-tag and TEV cleavage site, and a C-terminal 6xHis tag (lane 2). Bacterial lysates were incubated with nickel resin, and proteins washed (lane 3) or eluted (lane 4) from the resin, and following incubation with TEV protease (lane 5), were resolved by SDS-PAGE and stained with Coomassie Blue. The arrow shows tag-cleaved BCP1 NTase, and the asterisk indicates TEV protease.

(B) Bacterial cell lysates prepared from postinduction cultures of bacteria expressing SBP-tagged BCP1 mutant NTase (lane 1) were incubated with streptavidin resin, and proteins eluted from the resin following treatment with TEV protease (lane 2) were resolved by SDS-PAGE and stained with Coomassie Blue.

(C) Total protein was recovered from uninduced bacteria (lane 1) or postinduction cultures of bacteria expressing SBP-tagged Av05 NTase (lane 2). Bacterial cell lysates were incubated with streptavidin resin, and proteins eluted from the resin following treatment with TEV protease (lane 3) were resolved by SDS-PAGE and stained with Coomassie Blue.

**Figure S2. *Cg*TIR-STING is refractory to inhibition by BCP1 phage NTase, Related to Figure 2**

A) TIR NADase assay. *Capnocytophage granulosa* (*Cg*) NAD+ cleavage activity was measured using the fluorescent substrate ε-NAD, in the absence or presence of increasing amounts of the BCP1 phage NTase across a 10-fold range in concentration. Bar graphs show mean values, error bars represent s.d. Data are shown for n=4 experiments.

(B) CFU assay showing relative viability of *E. coli* transformed with the indicated combinations of *Cg*TIR-STING and wild-type BCP1 NTase or vector control. Results are representative of n=2 experiments.

**Figure S3. Purification of Atd1 and pyrophosphatase assay, Related to Figure 3**

(A) Total protein was recovered from uninduced bacteria (lane 1) or postinduction cultures of bacteria expressing Atd1 containing an N-terminal TS-tag and TEV cleavage site, and a C-terminal 6xHis tag (lane 2). Bacterial lysates were incubated with nickel resin, and proteins eluted from the resin (lane 3) were resolved by SDS-PAGE and stained with Coomassie Blue.

(B) Pyrophospatase assay measuring enzymatic activity of Atd1 on rNTP and cGG substrates. Results from two independent replicate experiments are plotted for each substrate.
